# An ancient satellite repeat controls gene expression and embryonic development in *Aedes aegypti* through a highly conserved piRNA

**DOI:** 10.1101/2020.01.15.907428

**Authors:** Rebecca Halbach, Pascal Miesen, Joep Joosten, Ezgi Taşköprü, Bas Pennings, Chantal B.F. Vogels, Sarah H. Merkling, Constantianus J. Koenraadt, Louis Lambrechts, Ronald P. van Rij

## Abstract

Tandem repeat elements such as the highly diverse class of satellite repeats occupy large parts of eukaryotic chromosomes. Most occur at (peri)centromeric and (sub)telomeric regions and have been implicated in chromosome organization, stabilization, and segregation^1^. Others are located more dispersed throughout the genome, but their functions remained largely enigmatic. Satellite repeats in euchromatic regions were hypothesized to regulate gene expression *in cis* by modulation of the local heterochromatin, or *in trans* via repeat-derived transcripts^2,3^. Yet, due to a lack of experimental models, gene regulatory potential of satellite repeats remains largely unexplored. Here we show that, in the vector mosquito *Aedes aegypti*, a satellite repeat promotes sequence-specific gene silencing via the expression of two abundant PIWI-interacting RNAs (piRNAs). Strikingly, whereas satellite repeats and piRNA sequences generally evolve extremely fast^4-6^, this locus was conserved for approximately 200 million years, suggesting a central function in mosquito biology. Tandem repeat-derived piRNA production commenced shortly after egg-laying and inactivation of the most abundant of the two piRNAs in early embryos resulted in an arrest of embryonic development. Transcriptional profiling in these embryos revealed the failure to degrade maternally provided transcripts that are normally cleared during maternal-to-zygotic transition. Our results reveal a novel mechanism in which satellite repeats regulate global gene expression *in trans* via piRNA-mediated gene silencing, which is fundamental to embryonic development. These findings highlight the regulatory potential of this enigmatic class of repeats.

## Main

Even though satellite repeats have been discovered nearly 60 years ago^7,8^, and comprise a substantial portion of eukaryotic genomes, little is known about the functions of this class of repetitive DNA. Many satellite repeats are actively transcribed, and some of them produce small interfering (si)RNAs required for the establishment and maintenance of heterochromatic regions^9-16^. Around two-thirds of the genome of *Aedes aegypti*, the most important vector for arthropod-borne viruses like dengue, Zika, and yellow fever virus, consists of repetitive elements^17^ (Extended Data Fig 1A), making this mosquito an interesting model to study these sequences. We analyzed small RNAs derived from unique and repetitive sequences in the genome of *Ae. aegypti* somatic and germline tissues as well as Aag2 cells. Even though satellite repeats constitute less than 10% of the genome, they were not only highly covered by siRNAs (Extended Data Fig 1A), but especially by PIWI-interacting (pi)RNAs (Extended Data Fig 1A). piRNAs are a class of small RNAs that protect animal genomes from harmful parasitic elements like transposons^18^. In the fruit fly *Drosophila melanogaster*, piRNAs are mostly derived from transposon-rich genomic regions termed piRNA clusters^19^. Yet, in *Ae. aegypti*, piRNAs from transposable elements (TEs) are underrepresented compared to their abundance in the genome^20^, especially in the soma (Extended Data Fig 1A), but instead, we found satellite-repeat derived piRNAs to be highly overrepresented in somatic tissues. Intriguingly, approximately three-quarters of these reads in the soma, and half of the reads in the germline or Aag2 cells represent only two individual sequences that map to a repeat locus on chromosome 3. This locus was about 3.5 kb in size and consisted of 20 full and one disrupted repeat unit organized in a head-to-tail array (Fig 1A, B). These two highly abundant satellite-derived small RNAs were 30 and 29 nucleotides in size, respectively (Extended Data Fig 1B), and resistant to β-elimination, suggesting that they are 2’-*O*-methylated at their 3’ end, a common feature of mature PIWI-bound piRNAs^21-23^ (Fig 1C). We named these two sequences tapiR1 and 2 (<u>t</u>andem repeat-<u>a</u>ssociated <u>piR</u>NA1/2). Expression of tapiR1 and 2 was ubiquitous in both somatic and germline tissues of adult mosquitoes (Extended Data Fig 2A). In *Ae. aegypti*, the PIWI-interacting RNA pathway has expanded to include seven PIWI genes (Piwi2-7 and Ago3) compared to three in flies^24^. Immunoprecipitation (IP) in Aag2 cells of the aedine PIWI proteins that are expressed both in the soma and gonads (Piwi4-6 and Ago3) followed by northern blotting or deep sequencing indicates that both tapiR1 and tapiR2 exclusively associate with Piwi4 (Fig 1D, Extended Data Fig 2B, C, Supplementary Fig S1A,B). Indeed, only knockdown of Piwi4, but not of other PIWI or AGO-clade genes reduced tapiR1 and 2 levels (Extended Figure 2D, E, Supplementary Fig S1C,D). Thus far, the piRNA repertoire and function of Piwi4 remained unclear. Piwi4 neither associates with TE nor virus-derived piRNAs^25^, yet was linked to piRNA biogenesis from transposons^25^ and to antiviral defense^26,27^. As nearly 90 % of Piwi4-associated small RNAs only comprise tapiR1, and, to a much lower extent, tapiR2 (Fig 1D), we hypothesize that tapiR1 not only dominates the piRNA repertoire, but also shapes downstream functions of Piwi4.

**Figure 1:**
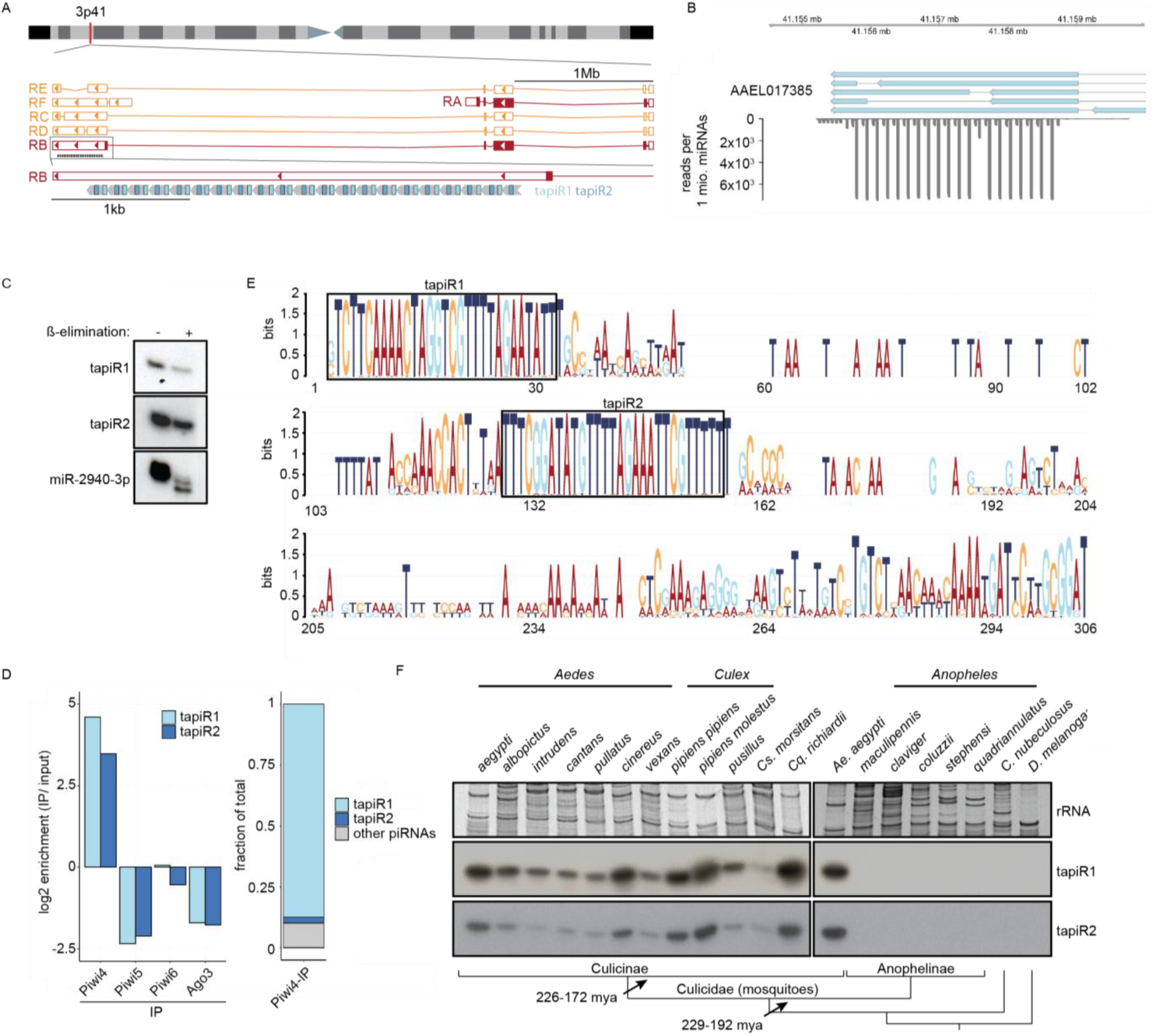
Conserved piRNAs are expressed from a satellite repeat and associate with Piwi4. (A) Current annotation of the gene AAEL017385 and its splice variants (RA-RF) on chromosome 3, with the position of the satellite repeat locus and tapiR1/2 piRNAs indicated. (B) Read coverage of the satellite repeats locus. Depicted are exons of AAELL017385 (blue), and small RNAs per million mapped miRNAs in Aag2 cells. (C) Small RNA nothern blot of tapiR1 and 2 upon ß-elimination in Aag2 cells. miR-2940-3p serves as positive control for the treatment. (D) Enrichment or depletion of tapiR1/2 compared to input sample in the indicated PIWI-IP small RNA sequencing libraries (left panel), and fraction of tapiR1/2 on total reads enriched in Piwi4 (right panel). (E) Sequence conservation of the satellite repeat monomers. All individual repeat monomers from *Ae. aegypti, Ae. albopictus* and *Cx. quinquefasciatus* were used to generate the sequence logo. Boxes highlight the position of tapiR1 and 2 in the monomer. (F) Northern blot analysis of tapiR1/2 in the indicated mosquito species (genera *Aedes, Culex, Culiseta, Coquillettidia*, and *Anopheles*) and other insects (*Culicoides* and *Drosophila*). Ethidium bromide-stained rRNA serves as loading control. For comparison, *Ae. aegypti* was included twice. Schematic representation of the phylogenetic relationships are indicated in the bottom panel. Bar lengths are arbitrary and do not reflect evolutionary distances.

**Figure 2:**
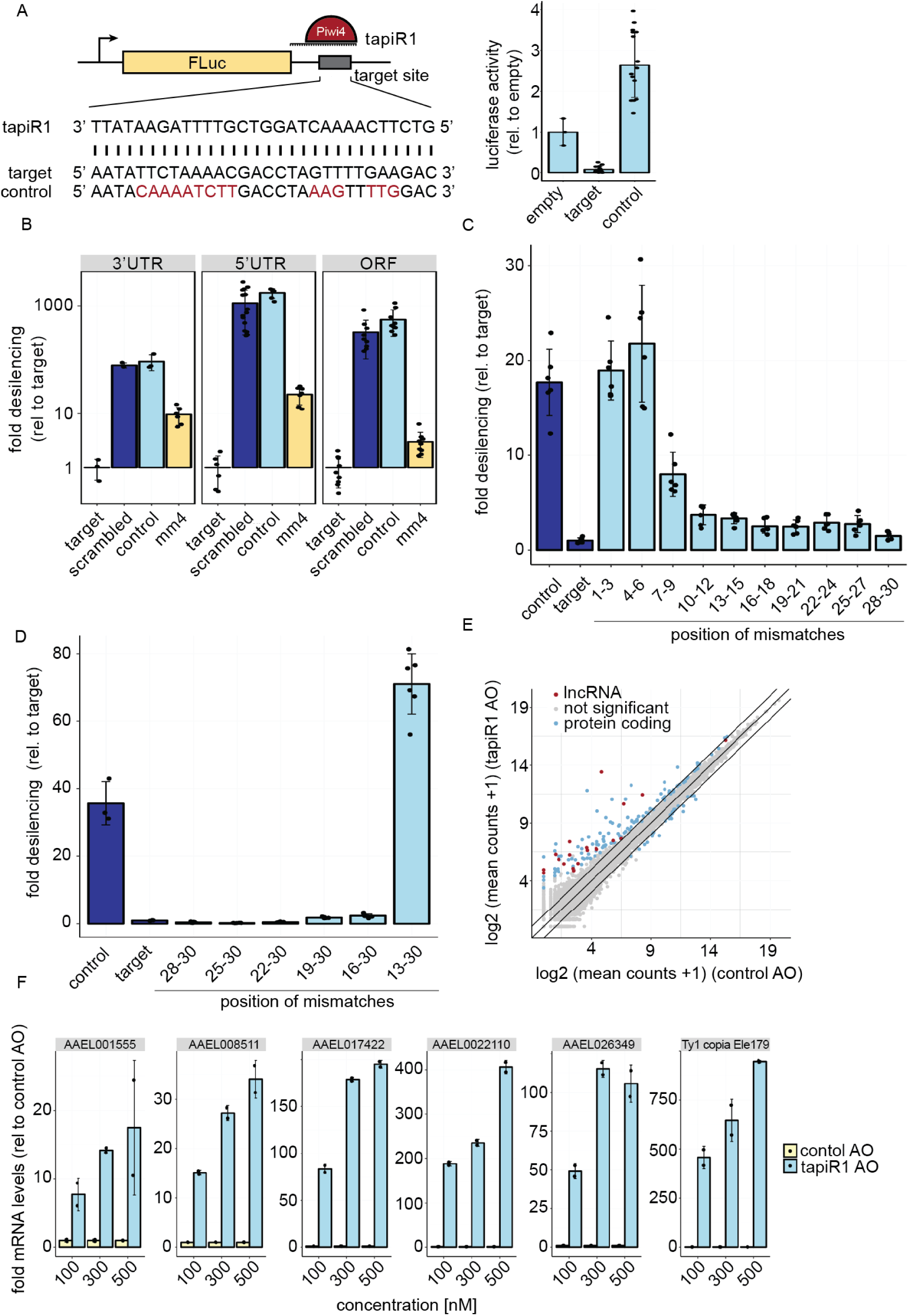
tapiR1 silences target RNAs *in trans* through seed-mediated base pairing. (A) Schematic representation of the firefly luciferase (FLuc) reporter constructs (left panel) and luciferase assay in Aag2 cells transfected with reporters containing no target site (empty), a fully complementary target site to tapiR1, or a control target site. (B) Luciferase assay of reporters with tapiR1 target sites, mismatched sites (mm4), or control sequences located at different positions in the reporter mRNA. (C, D) Luciferase assay of tapiR1 reporters harbouring three consecutive mismatches (C), or increasing number of mismatches (D) at the indicated positions of the piRNA target site in the 3’ UTR of firefly luciferase. Firefly luciferase activity was normalized to the activity of a co-transfected *Renilla* luciferase reporter. Indicated are mean, standard deviation, and individual measurements of a representative experiment performed with two to three independent clones per construct and measured in triplicates. (E) log2 expression of mRNAs and lncRNAs in Aag2 cells upon treatment with a tapiR1 specific or control antisense oligonucleotide (AO). Depicted are average read counts in three biological replicates. A pseudo-count of one was added to all values in order to plot values of zero. Diagonal lines indicate a fold change of two. Significance was tested at a false discovery rate (FDR) of 0.01 and a log2 fold change of 0.5 as indicated by coloured dots. (F) RT-qPCR of tapiR1 target genes upon transfection of Aag2 cells with tapiR1 specific or control AO. Depicted are mean, standard deviation, and individual measurements of one experiment measured in technical duplicates.

Satellite repeats are one of the fastest evolving parts of eukaryotic gnomes. Except for a few examples^28-30^, most satellite repeats display high sequence divergence between species, and can even be species-specific^6,31,32^, akin to piRNAs^4,5^. Hence, we were surprised to find that the identified tandem repeat locus in *Ae. aegypti* is conserved in the closely related Asian tiger mosquito *Ae. albopictus*, and even in the more distantly related southern house mosquito *Culex quinquefasciatus* (Extended Data Fig 3A). This locus is, however, not present in the genome assembly of the malaria vector *Anopheles gambiae*. The tandem repeat locus differed in the number of monomers across species, and the monomers exhibited substantial length and sequence divergence, both between species and between monomers within one species. However, the parts of the monomer that give rise to tapiR1 and 2 were by far more conserved than the overall monomer, suggesting that these sequences are under extensive selective constraints (Fig 1E). We further analyzed whether expression of the two repeat locus-derived piRNAs is conserved among different mosquitoes, including species for which no genome assembly is available. We analyzed 17 different mosquito species from 5 different genera (*Aedes, Culex, Culiseta, Coquillettidia* and *Anopheles*), as well as *Culicoides nubeculosus*, a biting midge that also transmits arboviruses, but is only distantly related to mosquitoes, and the fruitfly *Drosophila melanogaster*. Strikingly, even though piRNAs are usually not conserved even between closely related species^4,5^, we detected both tapiR1 and 2 in four genera of the Culicinae subfamily of mosquitoes (Fig 1F, Extended Data Fig 3B). In line with the absence of the repeat locus in the *Anopheles gambiae* genome, we did not observe tapiR1 or 2 expression in this subfamily of mosquitoes, nor in the two non-mosquito species. This observation suggests that the locus evolved in the late Triassic after divergence of the Anophelinae from the Culicinae subfamily of mosquitoes (229-192 mya^33^), but before further divergence of the culicine genera (226-172 mya^33^). This establishes this repeat locus as one of the very few ancient and deeply conserved satellite repeats that have hitherto been described^28-30,34^. Conservation of the locus over million years of mosquito evolution strongly suggests important and conserved functions for the locus and its associated piRNAs.

The satellite repeat locus overlaps with the 3’UTR of the gene AAEL017385 (LOC23687805) of unknown function in the current genome annotation (Fig 1A). This organization suggests that one or more splice variants of AAEL017385 are the source of the piRNAs, and that expression of this gene and piRNA biogenesis might be closely linked. However, knockdown of the different splice isoforms did not reduce expression of tapiR1 (Extended Data Fig 3C, Supplementary Fig S1E), arguing against this possibility. Rapid amplification of 3’ ends (RACE) of AAEL017385 transcripts revealed transcription termination sites directly upstream of the first tapiR1 and tapiR2 repeat, respectively (Extended Data Fig 3D). Even though we cannot exclude that some AAEL017385 transcripts overlap with the satellite repeat, our data strongly suggest that the two piRNAs and AAEL017385 are not expressed from the same transcriptional unit. Instead, the satellite repeat locus might be transcribed from an unknown upstream or internal promoter. In support of this notion, the repeat locus is not associated with the AAEL017385 orthologues in *Ae. albopictus* (AALF011179) and *Cx. quinquefasciatus* (CPIJ011773).

We next characterized the sequence requirements for target silencing by tapiR1, as its expression is approximately one log higher compared to tapiR2 in Aag2 cells. Using a luciferase reporter harbouring a fully complementary target site in the 3’ UTR, we validated that this piRNA is able to target RNAs *in trans*. The reporter was silenced more than 10-, or 35-fold compared to a reporter without target site, or a control reporter with a partially inverted target site, respectively (Fig 2A). Addition of an antisense oligonucleotide (AO) complementary to tapiR1 relieved silencing in a concentration-dependent manner (Extended Data Fig 4), confirming that the observed effect is mediated by the piRNA in a sequence-specific fashion. Unlike most miRNAs^35^, silencing was not dependent on the position of the target site in the mRNA and was efficient in the open reading frame and both the 5’ and the 3’ UTR (Fig 2B). During the course of our study we noticed that the firefly and *Renilla* luciferase genes, which we used as reporter and normalization control, respectively, contain potential target sites for tapiR1. We confirmed that *Renilla* luciferase is indeed potently suppressed by tapiR1 and firefly luciferase slightly (Extended Data Fig 5A-C), yet, mutating these target sites did not affect any of the conclusions reported below, but increased the observed effects (Extended Data Fig 5D,E).

To assess targeting requirements for tapiR1, we introduced mismatches in the piRNA target sites. Three consecutive mismatches were tolerated unless they were located in the t1 to t9 region of the piRNA (the nucleotides based-paired to piRNA positions 1 to 9) (Fig 2C, Extended Data Fig 6A), and single mismatches only impaired silencing at positions t3 to t7 (Extended data Fig 6A,B), reminiscent of a microRNA seed^35^ and comparable to what has been termed the piRNA seed in *C. elegans*^*36,37*^. Even though tapiR1 targeting requirements resemble those of miRNAs, the results are unlikely to be due to the piRNA being funnelled into the miRNA pathway. First, tapiR1 biogenesis is not dependent on Ago1 (Extended Data Fig 2E), and secondly, this piRNA is 2’-*O*-methylated (Fig 1C), a feature of siRNAs and mature PIWI-bound piRNAs, but not Ago1-associated miRNAs. A mismatch at position t1 did not alter silencing, suggesting that the first nucleotide is anchored in a binding pocket of Piwi4, similar to other Argonaute proteins^38-40^. Unexpectedly, a mismatch at position t2 was tolerated as well. This was, however, only the case when the rest of the target site was perfectly complementary, but not when the target contained mismatches outside of the seed (Extended Data Fig 6C). We further noticed that, in contrast to *C. elegans*^36^, G:U wobble pairs were not tolerated inside the seed and had the same effect as a mismatch at the same position (Extended Data Fig 6D). Whereas the 5’ seed region is normally absolutely required for targeting^36-38,41,42^, the 3’ part of the piRNA might increase specificity and efficiency of the targeting. For this reason we further assessed the extent of complementarity needed to allow for tapiR1-mediated silencing. Introduction of increasing numbers of mismatches at the 3’ end did not interfere with silencing when at least half of the piRNA could base pair with the target site (Fig 2D), indicating that the 3’ part of the piRNA is not necessarily required, yet the seed region alone not sufficient for silencing.

Taken together, these results suggest that tapiR1 needs relatively low sequence complementarity to efficiently silence targets^36,41-43^, and that there are no constraints regarding the position of the target site on the mRNA. As a consequence, tapiR1 may target a plethora of different cellular RNAs that are perfectly base pairing to the seed and additional matches outside the seed.

Considering the fact that the repeat locus is extremely conserved, we hypothesized that this piRNA regulates cellular gene expression. Some satellite repeats can influence genes by modulation of the local chromatin environment *in cis* ^14,44^, or have been hypothesized to induce silencing of genes with homologous repeat insertions^29^. In contrast, the tapiR1/2 locus has the potential to silence expression of a broad range of remote genes *in trans*, independent of repeat insertions in target genes, and thus, to regulate diverse and highly complex cellular processes.

To test this idea, we blocked tapiR1-mediated silencing with the tapiR1 AO in Aag2 cells and assessed global gene expression by RNAseq two days after treatment. Intriguingly, expression of 134 genes, amongst which many long non-coding RNAs, was significantly increased up to around 450 fold compared to the treatment with a control oligonucleotide (Fig 2E, Supplementary Table 1). Transposons were not globally affected, although some elements were up-regulated as well, up to around 850-fold (Extended Data Fig 7A, Supplementary Table 2). Expression of deregulated genes, and a transposable element was increased in a concentration-dependent manner upon tapiR1 AO treatment as measured by RT-qPCR (Fig 2F), validating our RNAseq results. We then used RNAHybrid to predict tapiR1 target sites and verified these sequences in luciferase reporter assays. Twelve out of 23 sites in protein-coding genes, lncRNAs, or transposable elements were indeed sufficient to support suppression of the reporter (Extended Data Fig 8), strongly suggesting that tapiR1 directly represses these cellular RNAs, and confirming that tapiR1 only needs limited base pairing to mediate silencing. Computational prediction of target sites has inherent limitations, and not all genes with predicted target sites were differentially regulated upon AO treatment, or, *vice versa*, target sites were functional in a reporter context, yet the gene itself was not differentially expressed (Extended Data Fig 7B,C). Additionally, the minimum free energy of the piRNA/target duplex was not predictive for the effect size of tapiR1-mediated silencing. Thus, similar to miRNAs^45^, other factors beyond Watson-Crick base pairing seem to play a role in definition of *bona fide* target genes for tapiR1. Nevertheless, our results indicate that tapiR1 is able to directly and strongly regulate gene expression and transposon RNA levels in a sequence-dependent manner. The expression of satellite repeats is often regulated in a developmental or stage-specific manner^29,46^, thus we analysed the expression pattern of tapiR1 and 2 throughout the mosquito life cycle. tapiR1 and 2 were not expressed during the first three hours of embryonic development of *Ae. aegypti* (Fig 3A), but could be detected in all subsequent life stages (Extended Data Fig 9A,B). At the very beginning of embryonic development the zygotic genome is transcriptionally quiescent and the first mitotic divisions are exclusively driven by maternally deposited transcripts and proteins. Maternal-to-zygotic transition (MZT) is marked by the degradation of these maternal transcripts and concomitant zygotic genome activation^47^. Destabilization of maternal transcripts initially occurs through maternal decay activities, and later, after onset of zygotic transcription also by zygotic components^47,48^. One of the best described mechanisms involves zygotically expressed miRNAs, for example miR-430, miR-427, and the miR-309 cluster in zebrafish^49^, *Xenopus*^50^, and flies^51^, respectively. Based on its expression pattern and its strong suppressive ability we hypothesized that tapiR1 could be part of the zygotic degradation pathway in mosquitoes, and necessary for embryonic development. To test this idea we injected either a tapiR1-specific AO or control AO into early pre-blastoderm *Ae. aegypti* embryos (before zygotic genome activation and expression of tapiR1) (Fig 3B), and assessed their development using discrete scoring schemes (Supplementary Fig 1F). Strikingly, more than 90 % of all tapiR1 AO-injected embryos were arrested early in development, whereas about half of all control embryos showed obvious signs of developmental progression (Fig 3C). In accordance, only a small fraction of tapiR1 AO-injected embryos hatched (Fig 3D) and continued to develop as larvae, suggesting that tapiR1-deficiency impedes development. RNA sequencing from tapiR1 AO or control AO-injected embryos 20.5 h after injection revealed massive deregulation of cellular transcripts (Fig 3E, Extended Data Fig 9C, Supplementary Table 3, 4). Expression of 205 genes, among which 44 lncRNAs, as well as few transposable elements was increased up to around 1000 and 500 fold in tapiR1 AO-treated embryos, respectively. Target genes with predicted tapiR1 target sites were more strongly up-regulated than genes without predicted site, as indicated by a shift of the cumulative distribution of RNA fold changes (Fig 3F). These findings show that tapiR1 controls regulatory circuits by direct gene targeting also *in vivo*, and that this function is essential for embryonic development, likely by promoting mRNA turnover of a subset of maternal transcripts. In line with this conclusion, confirmed target genes are down-regulated after the onset of tapiR1 expression (Fig 3G), and tapiR1 targets are overrepresented in transcripts that are maternally provided and degraded during MZT (Fig 3H).

**Figure 3:**
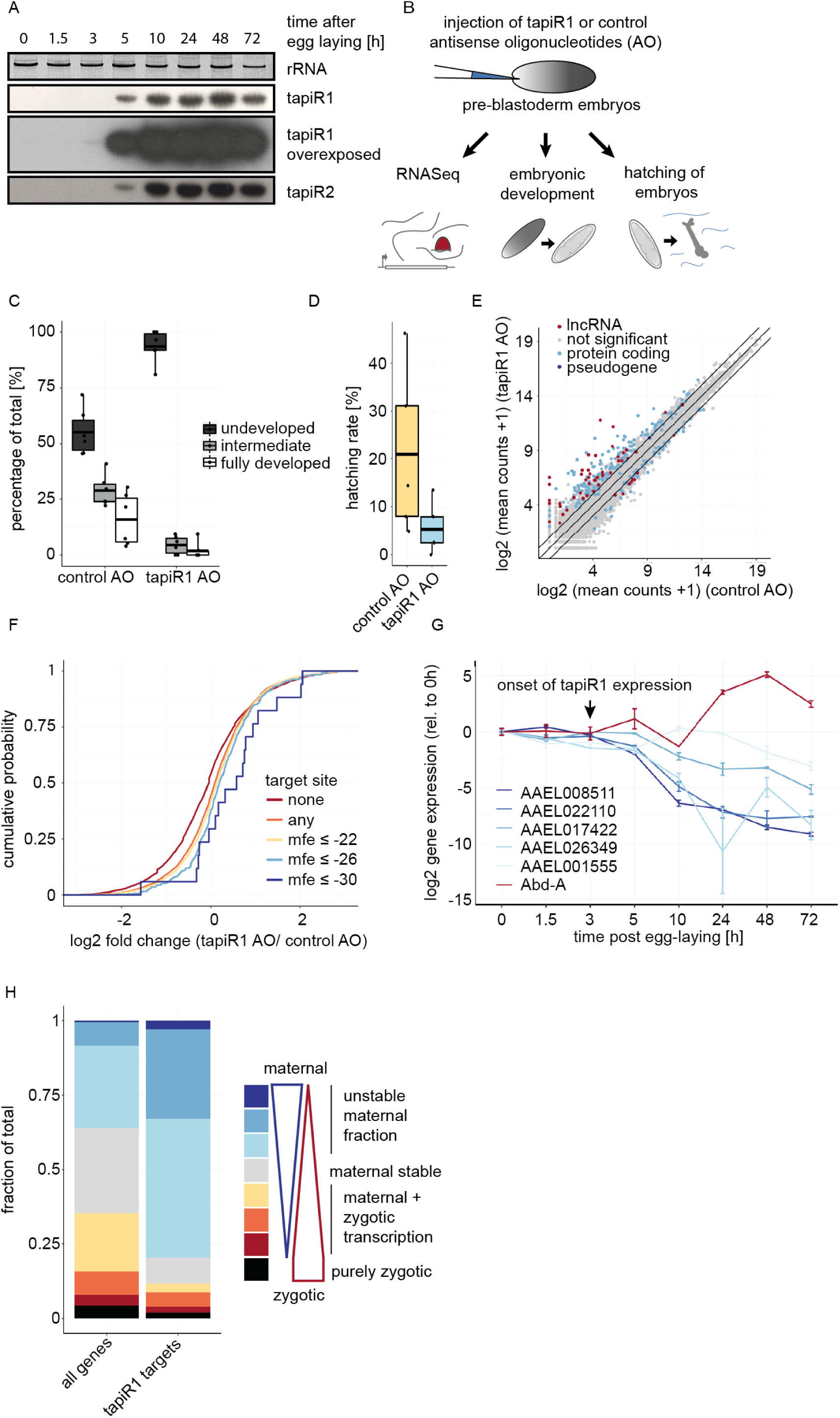
tapiR1 is essential for embryonic development *in vivo* by promoting turnover of maternally deposited transcripts. (A) Expression of tapiR1 and 2 as analysed by northern blot in *Ae. aegypti* embryos. Time indicates the age of the embryos after a 30 min egg laying period. For each time point, around 50 to 150 eggs were pooled. (B) Outline of the experimental procedure. (C) Percent of embryos injected with either tapiR1 or control AO that reached the indicated developmental stages 2.5 days post injection. Individual embryos were scored as either undeveloped, intermediate or fully developed, as shown in Supplementary Figure S1F. (D) Percent of embryos injected with tapiR1 or control AO that hatched four days post injection. Box-whiskers plot represents mean, first and third quartile and maximum and minimum of the data. Points show the individual experiments with 20 to 60 (C), or 50 to 150 (D) embryos per group. (E) log2 expression of genes in embryos injected with tapiR1 specific or a control AO at 20.5 h post injection. Mean counts from five biological replicates plus a pseudo-count of one are plotted. Per replicate, 50 embryos per group were pooled. Significance was tested at an FDR of 0.01 and a log2 fold change of 0.5. Diagonal lines highlight a fold change of two. (F) Experimental cumulative distribution of log2 fold changes of genes without or with predicted target sites for tapiR1. Target sites were grouped based on the predicted minimum free energy of the piRNA/target duplex. (G) Expression of tapiR1 target genes in *Ae. aegypti* embryos. RT-qPCR was performed on samples shown in (A). Abd-A is a gene not targeted by tapiR1 and serves as negative control. (H) Fraction of genes in different classes of genes expressed in embryos between 0 and 16 h post egg-laying.

piRNAs have been shown to promote degradation of *nanos*^52^ and other transcripts involved in germ cell development^53^. Yet, this was dependent on transposon-derived piRNAs, and rather depicts a re-purposing of the existing piRNA pool, which is, due to its large targeting potential, ideal to be used to degrade a large number of transcripts. In contrast, we propose that, analogous to abundant miRNAs in other animals^49-51^, Culicinae mosquitoes have evolved a specific piRNA to destabilize a defined set of maternally deposited transcripts in early embryonic development. To our knowledge, this is the first example of sequence-specific gene silencing by transcriptional products from a satellite repeat *in trans*, and underlines the regulatory potential of tandemly repeated DNA.

## Methods

### Cell culture

*Aedes aegypti* Aag2 cells were cultured in Leibovitz’s L-15 medium (Invitrogen) supplemented with 10 % heat-inactivated Fetal Bovine Serum (PAA Laboratories), 2 % Tryptose Phosphate Broth Solution (Sigma Aldrich), 1x MEM Non-Essential Amino Acids (Invitrogen) and 50 U/ml penicillin/streptomycin (Invitrogen) at 25 °C.

### Mosquito rearing

Injection and northern blot of embryos was performed using a Cell fusion agent virus-free, isofemale *Aedes aegypti* strain called Jane. This strain was initiated from a field population originally sampled in the Muang District of Kamphaeng Phet Province, Thailand^54^, and reared for 26 generations at 28 ±1°C, 75±5 % relative humidity, 12:12 hour light-dark cycle. Embryos were hatched under low pressure for 30-60 min. Larvae were grown in dechlorinated tap water and fed fish food powder (Tetramin) every two days. Adults were maintained in cages with constant access to a 10% sucrose solution. Female mosquitoes were fed on commercial rabbit blood (BCL) through a membrane feeding system (Hemotek Ltd.) using pig intestine as membrane. For AO injections, female mosquitoes were transferred to 25 °C and 70 % humidity for at least two days before forced to lay eggs, and embryos were then placed back to 28 °C immediately after the injection. For the time-course experiment in Fig. 3A,G, embryos were kept at 25 °C during the course of the experiment.

All other *in vivo* experiments were performed with the *Ae. aegypti* Rockefeller strain, obtained from Bayer AG, Monheim, Germany. The mosquitoes were maintained at 27±°C with 12:12 hour light:dark cycle and 70% relative humidity, as described before^55^.

Mosquitoes used in Fig 1F were either different laboratory-reared, or wild-caught species: *Aedes aegypti* Liverpool strain, *Culex pipiens, Anopheles coluzzii, An. quadriannulatus, An. stephensi* mosquitoes, *Culicoides nubeculosus* biting midges, and *D. melanogaster* w^1118^ flies were laboratory strains. The mosquitoes were deep-frozen and stored at −80 °C until use. *Ae. albopictus, Ae. cantans, Ae. intrudens, Ae. pullatus, Ae. cinereus, Ae. vexans, Cx. pusillus, Culiseta morsitans, Coquillettidia richiardii, An. maculipennis, An. claviger*, and *An. coluzzii* were wild-caught individuals collected in different regions in Italy, Sweden, or the Netherlands between July 2014 and June 2015^56^. Species were identified at the species level, and stored at −20 °C for a maximum of two years.

### Gene knockdown

Double-stranded RNA was generated by *in vitro* transcription of T7 promoter-flanked PCR products with T7 RNA polymerase. Primer sequences are given in Supplementary Table S5. The reaction was carried out at 37°C for 3 to 4 h, then heated to 80 °C for 10 min and gradually cooled down to room temperature to facilitate dsRNA formation. The dsRNA was purified with the GeneElute Total RNA Miniprep Kit (Sigma Aldrich).

Aag2 cells were seeded in 24-well plates the day before the experiment, and transfected with X-tremeGENE HP Transfection reagent (Roche) according to the manufacturer’s instructions, using a ratio of 4 μL reagent per μg of dsRNA. The transfection medium was replaced after 3 h with fully supplemented Leibovitz-15 medium and RNA was harvested 48 h later. Knockdown was confirmed by RT-qPCR.

### RNA isolation

RNA from cells and mosquitoes was isolated with Isol-RNA lysis buffer (5PRIME) according to the manufacturer’s instructions. Briefly, 200 μL chloroform was added to 1 mL lysis buffer, and centrifuged at 16,060 x g for 20 min at 4 °C. Isopropanol was added to the aqueous phase, followed by incubation on ice for at least one hour, and centrifugation at 16,060 x g for 10 min at 4 °C. The pellet was washed three to five times with 85 % ethanol and dissolved in RNase free water. RNA was quantified on a Nanodrop photospectrometer.

### Periodate treatment and β-elimination

Total RNA was treated with 25 mM NaIO_4_ in a final concentration of 60 mM borax and 60 mM boric acid (pH 8.6) for 30 min at room temperature. In the control, NaIO_4_ was replaced by an equal volume of water. The reaction was quenched with glycerol and β-elimination was induced with a final concentration of 40 mM NaCl for 90 min at 45 °C. RNA was ethanol precipitated and analysed with northern blot.

### Generation of antibodies

Custom-made antibodies (Eurogentec) against endogenous PIWI proteins were generated by immunization of two rabbits per antibody with a mix of two unique peptides (Ago3: TSGADSSESDDKQSS, IIYKRKQRMSENIQF; Piwi4: HEGRGSPSSRPAYSS, HHRESSAGGRERSGN; Piwi5: DIVRSRPLDSKVVKQ, CANQGGNWRDNYKRAI; Piwi6: MADNPQEGSSGGRIR, RGDHRQKPYDRPEQS). After 87 days and a total of four immunizations (t=0, 14, 28, 56 days), sera of both rabbits were collected, pooled, and purified against each peptide separately. Specificity of the antibody was confirmed by Western blotting of Aag2 cells stably expressing PTH (ProteinA, TEV cleavage site, 6x His-tag)-tagged PIWI^57^ upon knockdown of the respective PIWI protein, or a control knockdown (dsRLuc) (see Supplementary Fig S1A).

### Immunoprecipitation and western blotting

Aag2 cells were lysed with RIPA buffer (10 mM Tris-HCl, 150 mM NaCl, 0.5 mM EDTA, 0.1 % SDS, 1 % Triton-X-100, 10 % DOC, 1x protease inhibitor cocktail), supplemented with 10 % glycerol and stored at −80 °C until use. The IP was performed with custom-made antibodies against Piwi4-6 and Ago3 (1:10 dilution) at 4 °C for 4 h on rotation. Protein A/G Plus beads (Santa Cruz) were added at a dilution of 1:10 and then incubated overnight at 4 °C on rotation. Beads were washed 3 times with RIPA buffer, and half was used for RNA isolation and protein analysis each. For RNA extraction, beads were treated with proteinase K for 2 h at 55 °C and isolated with phenol-chloroform extraction. Equal amounts of RNA for input and IPs were then analysed by northern blotting. For western blotting, the IP samples were boiled in 2x Laemmli buffer for 10 min at 95 °C, separated on 7.5 % SDS-polyacrylamide gels, and blotted on 0.2 μm nitrocellulose membranes (Bio-Rad) in a wet blot chamber on ice. Membranes were blocked for 1 h with 5 % milk in PBS-T (137 mM NaCl, 12 mM phosphate, 2.7 mM KCl, pH 7.4, 0.1 % (v/v) Tween 20) and incubated with PIWI-specific (dilution 1:1000) and Tubulin primary antibodies (rat anti-Tubulin alpha, MCA78G, 1:1000, Sanbio) overnight at 4 °C. The next day, membranes were washed three times with PBS-T and incubated with secondary antibodies conjugated to a fluorescence dye (IRDye 800CW conjugated goat anti rabbit, 1:10,000, Li-Cor, and IRDye 680LT conjugated goat anti rat, 1:10,000, Li-Cor) for 1 h at room temperature in the dark. After washing three times in PBS-T, signal was detected with the Odyssey-CLx Imaging system (Li-Cor).

### Northern blot

piRNAs were detected by northern blot analyses, as published in ref.^58^. Briefly, RNA was denatured at 80 °C for 2 min in Gel Loading Buffer II (Ambion) and size-separated on 0.5 x TBE (45 mM Tris-borate, 1 mM EDTA), 7 M Urea, 15 % denaturing polyacrylamide gels. RNA was then blotted on Hybond-NX nylon membranes (GE Healthcare) in a semi-dry blotting chamber for 45 min at 20 V and 4 °C and crosslinked to the membrane with EDC crosslinking solution (127 mM 1-methylimidazole (Sigma-Aldrich), 163 mM N-(3-dimethylaminopropyl)-N&-ethylcarbodiimide hydrochloride (Sigma-Aldrich), pH 8.0) at 60 °C for two hours. Crosslinked membranes were pre-hybridized in ULTRAHyb-Oligo hybridization buffer (Thermo Scientific) for one hour at 42 °C and probed with indicated ^32^P 5’ end-labelled DNA oligonucleotide probes over night at 42 °C. Membranes were then washed with decreasing concentrations of SCC (300 mM NaCl, 30 mM sodium citrate pH 7.0; 150 mM NaCl, 15 mM sodium citrate; 15 mM NaCl, 1.5 mM sodium citrate) and 0.1 % SDS, and exposed to Carestream BioMax XAR X-Ray films (Kodak). Probe sequences can be found in the Supplementary Table S5.

### Reporter cloning and luciferase assay

Reporters were constructed by cloning annealed and phosphorylated oligonucleotides with the indicated tapiR1 or control target sites in the pMT-GL3 vector^59^. This vector encodes the *Photinus pyralis* firefly luciferase (GL3) under a copper-inducible metallothionein promoter. Sense and antisense oligonucleotides (Sigma Aldrich) were annealed by heating to 80 °C, and gradually cooling down to room temperature, phosphorylated with T4 polynucleotide kinase (Roche) at 37 °C for 30 min, purified and then ligated into the pMT-GL3 vector. For cloning of 3’UTR and 5’ UTR reporters, the target site or the target site and an upstream *BamH*I site were cloned into the *Pme*I and *Sac*II, or *Not*I and *Xho*I restriction sites, respectively. ORF reporters were constructed by cloning a Kozak sequence followed by the first 45 nucleotides of luciferase and the target site into *Xho*I and *Nco*I sites. Sequences of the oligonucleotides are provided in Supplementary Table S5. Where indicated, mutated firefly or *Renilla* luciferase versions were used that harbour synonymous mutations destroying the predicted target sites for tapiR1 (firefly luciferase: 782 gagtcgtcttaatgtatagatttgaagaa 810 mutated to 782 g**t**gtcgt**gc**t**t**atgta**cc**g**g**tt**c**ga**g**ga**g** 810, and *Renilla* luciferase 462 tgaatggcctgatattgaagaa 483 mutated to 462 tga**g**tggcc**a**gatat**c**ga**g**ga**g** 483; modified nucleotides in bold).

Aag2 cells were seeded in 96-well plates the day before the experiment and transfected with 100 ng of the indicated plasmids and 100 ng pMT-*Renilla*^*59*^ per well, using 2 μL X-tremeGENE HP DNA transfection reagent per 1 μg plasmid DNA according to the manufacturer’s instructions. Alternatively, 100 ng reporter plasmid and 100 ng pMT-*Renilla* were co-transfected with the indicated amounts of unlabelled, fully 2’*O*-methylated antisense RNA oligonucleotide using an additional amount of 4 μL X-tremeGENE HP DNA transfection reagent (Roche) per 1 μg oligonucleotide. Medium was replaced 3 h after reporter plasmid transfection with 0.5 mM CuSO_4_ in fully supplemented Leibovitz’s L-15 medium to induce the metallothionein promoter. 24 h later, cells were lysed in 30 μL Passive lysis buffer (Promega) and activity of both luciferases was measured in 10 μL of the sample with the Dual Luciferase Reporter Assay system (Promega) on a Modulus Single Tube Reader (Turner Biosystems). Firefly luciferase was normalized to *Renilla* luciferase activity. For each construct, at least two to three independent clones were measured in triplicate.

### RT-qPCR

1 μg of total RNA was treated with DNaseI (Ambion) for 45 min at 37 °C and reverse transcribed using the Taqman reverse transcription kit (Applied Biosystems) according to the manufacturer’s protocol. Real-time PCR was performed with the GoTag qPCR Master Mix (Promega) and measured on a LightCycler480 instrument (Roche) with 5 min initial denaturation and 45 cycles of 5 s denaturation at 95 °C, 10 s annealing at 60 °C and 20 s amplification at 72 °C. Starting fluorescence values of specific mRNAs were calculated with linear regression method of log fluorescence per cycle number and LinRegPCR program, version 2015.3, as described in ref ^60^.

### 3’ RACE

3’ Rapid Amplification of cDNA Ends (3’ RACE) was performed using the FirstChoice RLM-RACE Kit (Thermo Fischer Scientific) according to the manufacturer’s instructions. Amplification products were separated on agarose gel, purified and Sanger sequenced. Primer sequences can be found in Supplementary Table S1.

### Blood feeding experiment

Naïve female *Aedes aegypti* (Liverpool strain) mosquitoes were offered human blood (Sanquin Blood Supply Foundation, Nijmegen, The Netherlands) through a Parafilm membrane using the Hemotek PS5 feeder (Discovery Workshops). Five engorged females were selected and sacrificed at each of the indicated time points. RNA was isolated as described above.

### tapiR1 antisense oligonucleotide treatment and injection

Aag2 cells were seeded in 24-well plates the day before the experiment. Cells were treated with 500 nM 5’Cy5-labelled, fully 2’*O*-methylated antisense RNA oligonucleotide in 530 μL medium with 4 μL X-tremeGENE HP DNA transfection reagent (Roche) per 1 μg oligonucleotide. Medium was replaced after 3 h and cells from three independent experiments were harvested 48 h after transfection and prepared for RNA sequencing (see below).

For injection of embryos, engorged female mosquitoes that were kept at 25 °C and 70 % humidity were allowed to lay eggs for 45 min. Embryos were desiccated for 1.5 min, covered with Halocarbon oil (Sigma Aldrich) and injected with 50 μM 5’Cy5-labelled, fully 2’*O*-methylated antisense RNA oligonucleotide with a FemtoJet 4x (Eppendorf) with 1200 hPa pressure. Injected embryos were then transferred to a wet Whatman paper and kept at 27 °C and 80 % humidity for the indicated times. Per experiment, 50 to 150 embryos were injected per condition.

### Scoring of embryo development and hatching

Injected embryos were allowed to develop for 2.5 days after injection on a moist Whatman paper and then fixed in 4% paraform aldehyde for 8 h to overnight. Afterwards, the pigment of the endochorion was bleached with Trpis solution^61^ (0.037 M sodium chlorite, 1.45 M acetic acid) for 24 to 48 h. Embryos were washed five times in PBS and images were taken with a EVOS FL imaging system (Thermo Fisher Scientific). Embryos with evident larval segmentation (head, fused thoracical elements and abdomen) were scored as fully developed and embryos without any evident structure of the ooplasm as undeveloped. Individuals that showed first signs of structural rearrangements of the ooplasm, but did not complete larval segmentation were scored as intermediate (see Supplementary Fig S1F). To avoid biases, the scoring was performed blindly. Hatching rate was counted from injected embryos 4 days post injection. Embryos were kept moist for two days and then allowed to slowly dry for the rest of the period. The embryos were transferred to water and then forced to hatch by applying negative pressure for a period of 30 min. The number of hatched L1 larvae was counted immediately afterwards.

### Sequence logo

Repeat monomers from the satellite repeat loci in *Ae. aegypti, Ae. albopictus*, and *Cx. quinquefasciatus* were extracted manually from the current genome annotations obtained from Vectorbase (*Aedes aegypti* Liverpool AaegL5, *Aedes albopictus* Foshan AaloF1, *Culex quinquefasciatus* Johannesburg CpipJ2). A repeat unit was defined as the sequence starting from the first tapiR1 nucleotide until one nucleotide upstream of the next tapiR1 sequence. Sequences were aligned using MAFFT (v7.397)^62^ (with options –genafpair –leavegappyregion --kimura 1 -- maxiterate 1000 --retree 1) and the sequence logo was constructed with the R package ggseqlogo ^63^.

### Small RNA sequencing

Small RNAs from Aag2 cells (input) or PIWI immunoprecipitations were cloned with the TruSeq small RNA sample preparation kit (Illumina) according to the manufacturer’s instructions. For the input sample, size selected 19-33 nt small RNAs purified from polyacrylamide gel were used to construct the library as described previously^64^, whereas IP samples were not extracted from gel.

Libraries were sequenced on an Illumina HiSeq 4000 instrument by Plateforme GenomEast (Strasbourg, France).

### mRNA sequencing

RNA was isolated from Aag2 cells 48 h after AO transfection (three independent experiments), or embryos 20.5 h after AO injection (50 embryos pooled per experiment from five independent experiments) with RNAsolv reagent following standard phenol-chloroform extraction (see above). Polyadenylated RNAs were extracted and sequencing libraries were prepared using the TruSeq stranded mRNA Library Prep kit (Illumina) following the manufacturer’s instructions, and sequenced on an Illumina Hi Seq 4000 instrument (2×50 bases).

### Analysis of mRNA sequencing

Reads were mapped to the *Ae. aegypti* genome AaegL5 as provided by VectorBase (https://www.vectorbase.org) with STAR (version 2.5.2b)^65^ in 2-pass mode: first mapping was done for all samples (options: --readFilesCommand zcat --outSAMtype None --outSAMattrIHstart 0 --outSAMstrandField intronMotif), identified splice junctions were combined (junctions located on the mitochondrial genome were filtered out, as these are likely false positives), and this list of junctions was used in a second round of mapping (with –sjdbFileChrStartEnd) and default parameters as above. Reads were quantified with the additional option –quantMode GeneCounts to quantified reads per gene. Alternatively, reads were quantified on TEfam transposon consensus sequences (https://tefam.biochem.vt.edu/tefam/get_fasta.php) with Salmon (v.0.8.2)^66^, default settings and libType set to “ISR”. Statistical and further downstream analyses were performed with DESeq2^67^ from Bioconductor. Significance was tested at an FDR of 0.01 and a log2 fold change of 0.5. tapiR1 target sites were predicted with the online tool from RNAHybrid^68^ with helix constraints from nucleotide two to seven, and no G:U wobble allowed in the seed. Predictions were made on the AaegL5.1 geneset as provided by VectorBase, and on TEfam transposon consensus sequences. For Fig4H, publicly available sequencing datasets^69^ (accession numbers: SRR923702, SRR923826, SRR923837, SRR923853, SRR923704) were mapped and quantified as described above. Genes were categorized on the basis of their expression in embryos at 0-2 h vs. 12-16 h post egg-laying. Genes not detected in the 0-2 h sample were defined as purely zygotic, and genes that did not increase or decrease by more than log_2_(0.5) as maternal stable. Genes that changed in expression by more than log_2_(0.5), log_2_(2), and log_2_(5) from 0-2 h to 12-16 h were categorized as maternal unstable fraction (decreased expression), or as genes that are maternally provided but are transcribed by the zygote as addition to the preloaded maternal pool (increased expression). tapiR1 targets were defined as genes that were significantly upregulated at least two fold in tapiR1 AO injected embryos and harbour a predicted tapiR1 target site (mfe <= −24). The code will be made available on GitHub upon publication.

### Analysis of small RNA sequencing

3’ sequencing adapters (TGGAATTCTCGGGTGCCAAGG) were trimmed from the sequence reads with Cutadapt (version 1.14)^70^ and trimmed reads were mapped with Bowtie (version 0.12.7)^71^ to the *Aedes aegypti* LVP_AGWG genome sequence AaegL5.1 obtained from VectorBase with at most 1 mismatch. Reads that mapped to rRNAs or tRNAs were excluded from the analyses. Alternatively, 3’sequencing adapters ((NNN)TGGAATTCTCGGGTGCCAAGGC) and three random bases were trimmed from publicly available datasets from *Ae. aegypti* somatic and germline tissues^72^ (SRR5961503, SRR5961504, SRR5961505, SRR5961506) and then processed as described above. Oxidized libraries, IPs and input sample were normalized to the total number of mapped reads, all other libraries to the total number of miRNAs (in millions). piRNAs that were at least two fold enriched in a PIWI-IP compared to the corresponding input sample and were present with at least 10 rpm in the IP sample were considered PIWI-bound. Mapping positions were overlapped with basefeatures and repeatfeatures retrieved from VectorBase and counted with bedtools^73^. Reads that mapped to two or more features were assigned to only one feature with the following hierarchy: open reading frames > non-coding RNAs (incl. lncRNAs, pseudogenes, snoRNAs, snRNAs, miRNAs) > LTR retrotransposons > Non-LTR retrotransposons (SINEs, LINEs, Penelope) > “Cut and paste” DNA transposons > other DNA transposons (Helitrons, MITEs) > satellite and tandem repeat features > DUST > other /unknown repeats. Accordingly, reads that mapped to a repeat feature and an intron or UTR were classified as repeat-derived, whereas all other reads mapping to introns or UTRs were considered as gene-derived. Positions not overlapping with any annotation were summarized as “other”. Results were then visualized with ggplot2^74^ or Gviz^75^ in R.

The code will be made available on GitHub upon publication.

## Data availability

Raw sequence data is deposited in the NCBI Sequence Read Archive under the BioProject number PRJNA482553.

## Supplementary Information

Supplementary Table S1: Differentially expressed genes upon tapiR1 AO treatment in Aag2 cells.

Supplementary Table S2: Differentially expressed transposable elements upon tapiR1 AO treatment in Aag2 cells.

Supplementary Table S3: Differentially expressed genes upon tapiR1 AO treatment in embryos.

Supplementary Table S4: Differentially expressed transposable elements upon tapiR1 AO treatment in embryos.

Supplementary Table S5: Oligonucleotide sequences used in this study.

## Acknowledgments

We thank past and current members of the Van Rij laboratory for discussions. We are grateful to Anna Beth Crist and Artem Baidaliuk for their help with mosquito rearing and embryo injections, and to Catherine Bourgouin and Nicolas Puchot for assistance with the microinjection apparatus. We thank Bas Dutilh for his support with analyzing target site enrichments, and Geert-Jan van Gemert for kindly providing *An. stephensi* mosquitoes. Tim Möhlmann is acknowledged for providing wild-caught mosquito samples.

This work is financially supported by a Consolidator Grant from the European Research Council under the European Union’s Seventh Framework Programme (grant number ERC CoG 615680) and a VICI grant from the Netherlands Organization for Scientific Research (grant number 016.VICI.170.090). A stay of R.H. at Pasteur Institute, Paris, France was supported by ERASMUS+.

## Author contributions

R.H., P.M., and R.P.v.R designed the experiments and analyzed the data. R.H. performed the computational analyses and most of the experiments, except for PIWI-IPs for small RNA sequencing (J.J. and E.T.), design and validation of PIWI antibodies (B.P.), and tissue isolations and blood feeding experiment (C.B.F.V. and C.J.K.). S.H.M. and L.L. helped with optimizing embryo injections. R.H. and R.P.v.R. wrote the paper. All authors read and contributed to the manuscript.

## Author information

The authors declare no competing financial interests.

---

**Extended Data Figure 1:**
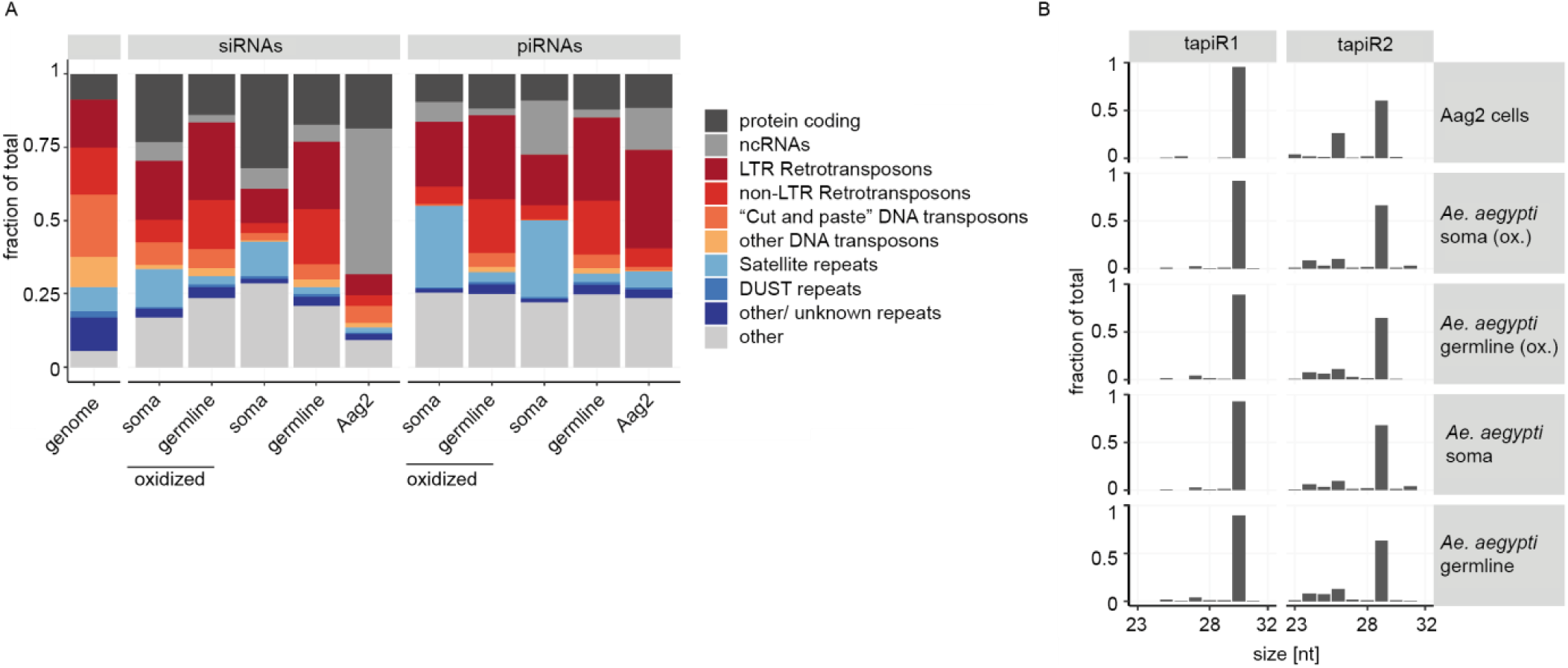
Expression of piRNAs from a satellite repeat locus. (A) Fraction of siRNAs and piRNAs mapping on genomic features in libraries derived from germline or somatic *Ae. aegypti* adult tissues. Small RNAs that overlapped multiple features were assigned to only one category (see Methods). Leftmost bar depicts the abundance of each feature category in the genome. (B) Read length distribution of tapiR1 and 2 in libraries from Aag2 cells, and adult germline and somatic tissues (oxidized or untreated).

**Extended Data Figure 2:**
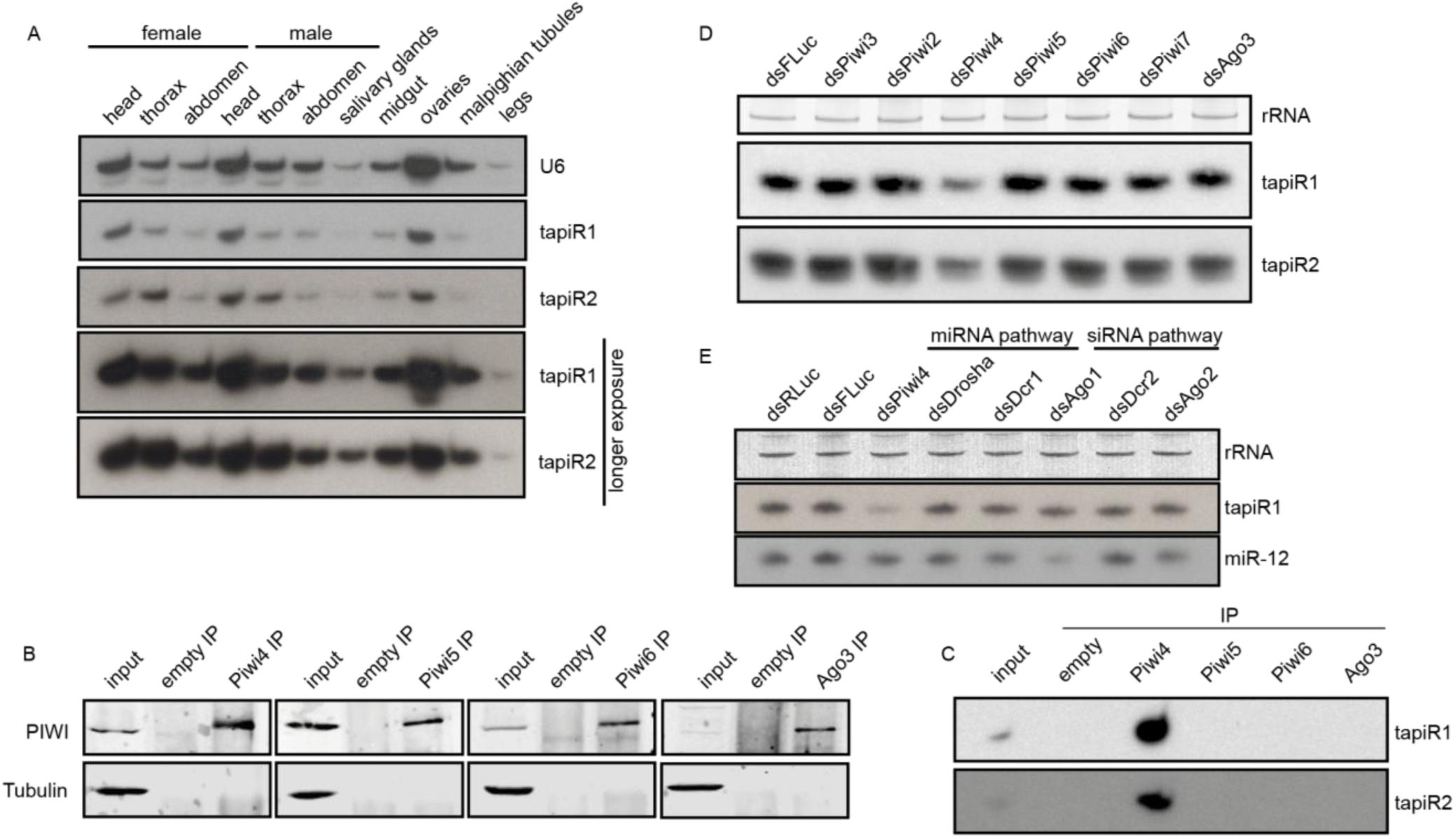
tapiR1 and 2 are expressed in *Ae. aegypti* mosquitoes and associate with PIWI proteins. (A, D, E) Northern blot of tapiR1 and 2 in different tissues of adult mosquitoes (A), upon dsRNA-mediated knockdown of individual PIWI genes (D), and upon knockdown of miRNA and siRNA pathway genes (E), or a control (dsFLuc and dsRLuc) in Aag2 cells. Ethidium bromide-stained rRNA, or U6 snRNA served as loading control. (B) Western blot analysis of the indicated PIWI proteins before (input) and after immunoprecipitation (IP) used for the small RNA northern blot of panel C. An IP with empty beads serves as negative control. Tubulin was used to control for non-specific binding. (C) Immunoprecipitation of the indicated PIWI proteins from Aag2 cells followed by northern blot analyses for tapiR1 and 2.

**Extended Data Figure 3:**
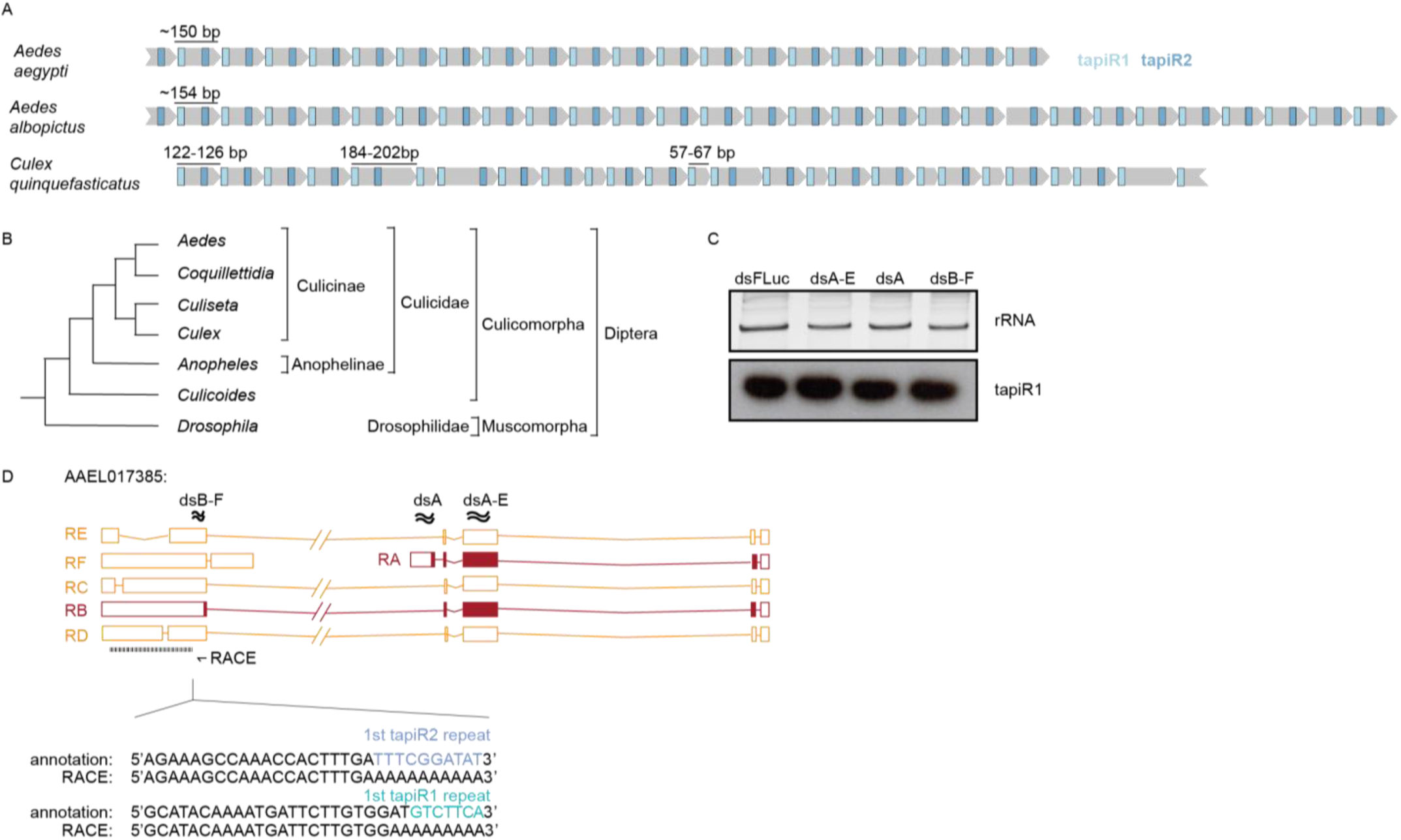
Expression of tapiR1 is independent of AAEL017385. (A) Schematic representation of the tapiR1/2 satellite repeat locus in *Ae. aegypti, Ae. albopictus* and *Cx. quinquefasciatus*. Numbers indicate lengths of the repeats, and, for *Cx. quinquefasciatus*, also length of deviating repeat monomers. (B) Evolutionary relationships of dipterous genera based on ref^33^. Bar lengths are arbitrary and do not reflect evolutionary distances. (C) Northern blot of tapiR1 in Aag2 cells treated with control dsRNA targeting different transcripts of AAEL017385, or, as control, firefly luciferase (FLuc). Ethidium bromide stained rRNA serves as loading control. (D) Top panel: Schematic representation of the AAEL017385 locus and satellite repeat. The primer used for 3’ RACE and positions targeted by dsRNA in panel C are indicated with an arrow and wavy lines, respectively. Bottom panel: 3’ RACE analysis of AAEL017385. Indicated are sequences from the current AaegL5 genome annotation and RACE PCR products. The sequences of the 5’ terminal tapiR1 and 2 repeats are highlighted with colours.

**Extended Data Figure 4:**
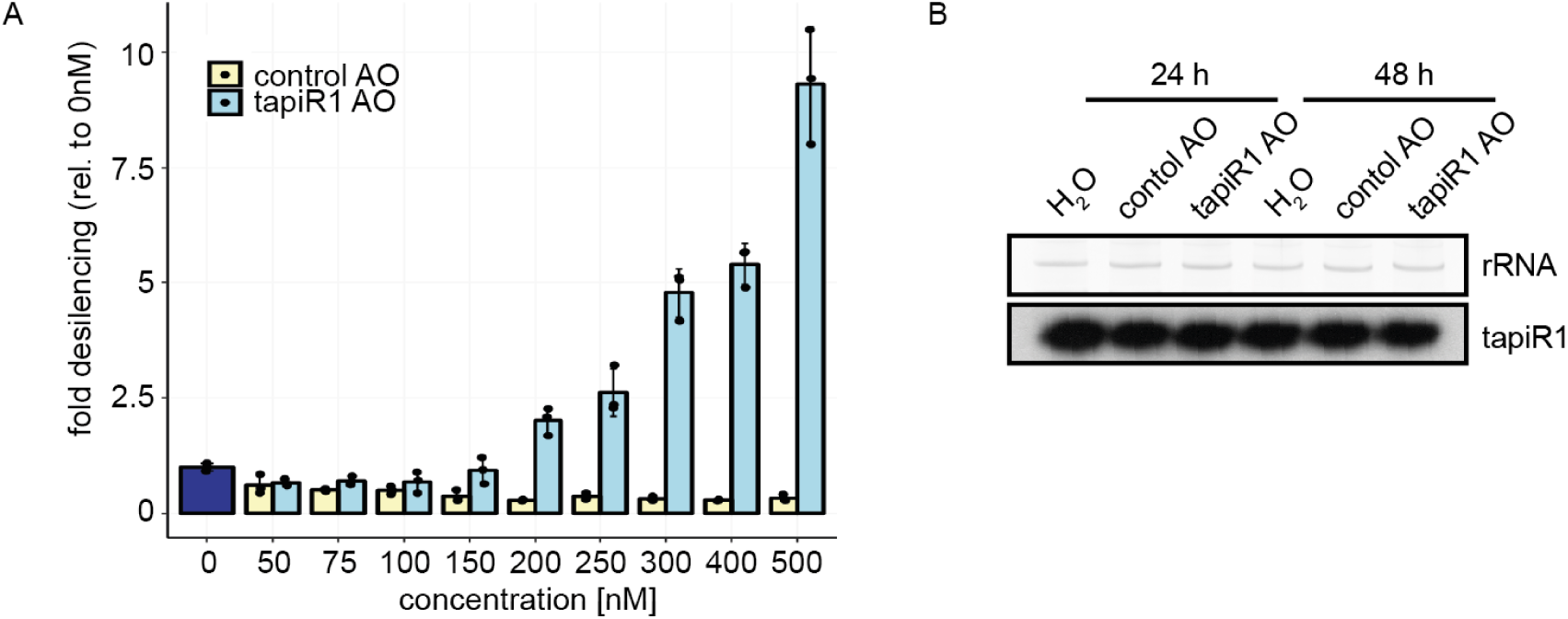
An antisense oligonucleotide relieves tapiR1-mediated silencing. (A) Luciferase assay of a reporter with a fully complementary target site for tapiR1 in the 3’ UTR. Cells were co-transfected with the reporter and increasing amounts of a fully 2’*O*-methylated antisense RNA oligonucleotide (AO), or a control AO. Firefly luciferase activity was normalized to the activity of a co-transfected *Renilla* luciferase reporter. Indicated are mean, standard deviation and individual measurements from a representative experiment measured in triplicate. (B) Northern blot detection of tapiR1 in Aag2 cells upon treatment with tapiR1 or control AO in Aag2 cells. Cells were harvested after the indicated time points. Ethidium bromide-stained rRNA serves as loading control.

**Extended Data Figure 5:**
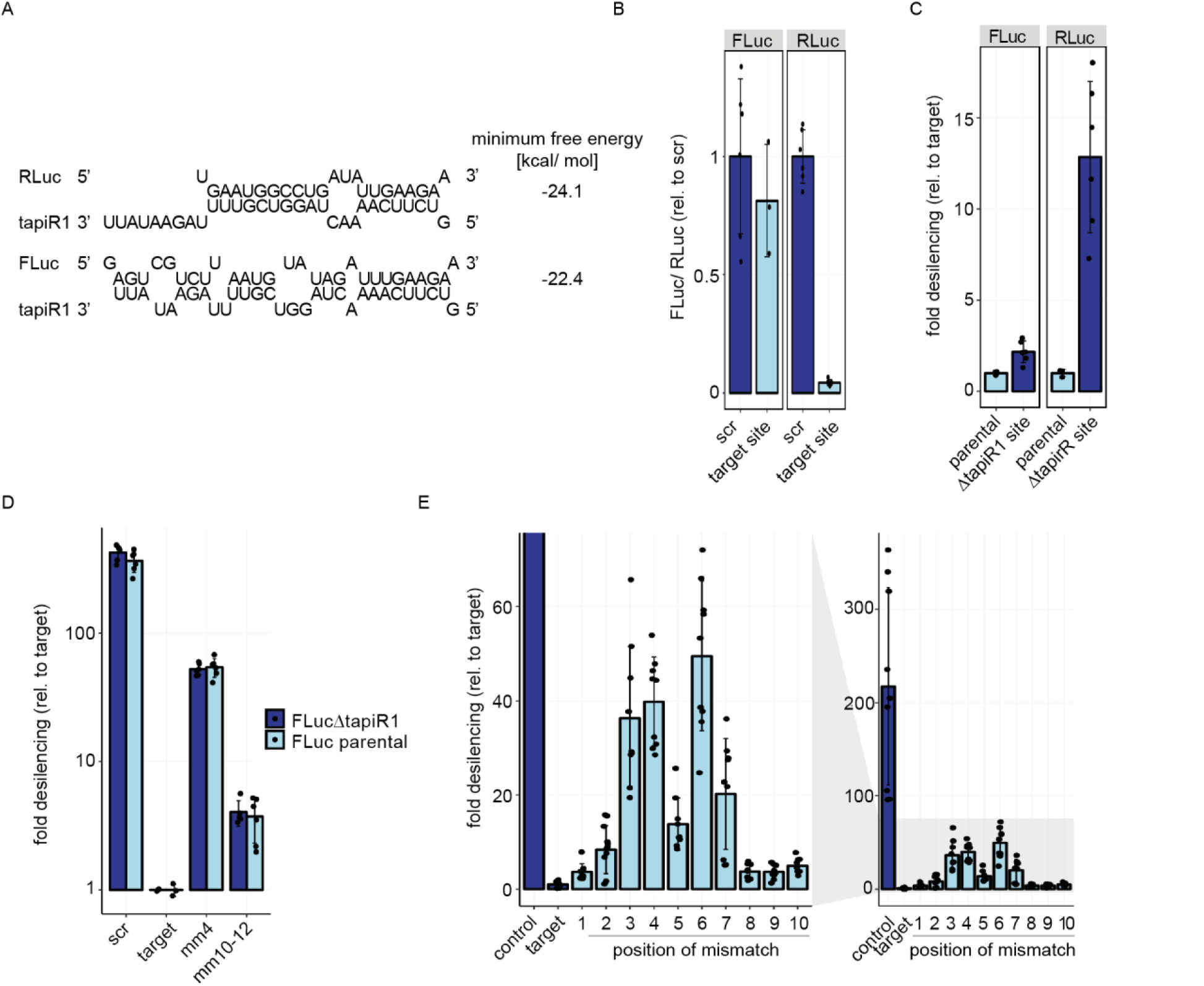
*Renilla* luciferase contains a functional tapiR1 target site. (A) Schematic representation of predicted tapiR1 target sites and minimum free energy of the indicated structures in the coding sequences of *Renilla* luciferase (RLuc) or firefly luciferase (FLuc). (B) Luciferase assay of Aag2 cells transfected with reporters carrying either a scrambled (scr) site or the predicted target site from firefly luciferase (left panel), or from *Renilla* luciferase (right panel) from panel (A) in the 3’UTR of FLuc. (C) Luciferase activity of firefly luciferase or *Renilla* luciferase construct with synonymous mutations introduced into the predicted tapiR1 target site (Δtapir1 site) or the parental clones. (D) Luciferase reporter assay of reporters carrying target sites for tapiR1 as indicated in panel A in the 3’UTR of either the parental firefly luciferase, or the ΔtapiR1 firefly luciferase version. (E) Reporter assay with luciferase carrying tapiR1 target sites with single mismatches in the 3’ UTR as used in Extended Data Fig 6B, using RLuc with a mutated tapiR1 target site (ΔtapiR1 site) for normalization. Left panel is a zoom to the x-axis of the right panel. Shown are mean, standard deviation and individual measurements from representative experiments performed with at least two different clones per construct, and measured in triplicate.

**Extended Data Figure 6:**
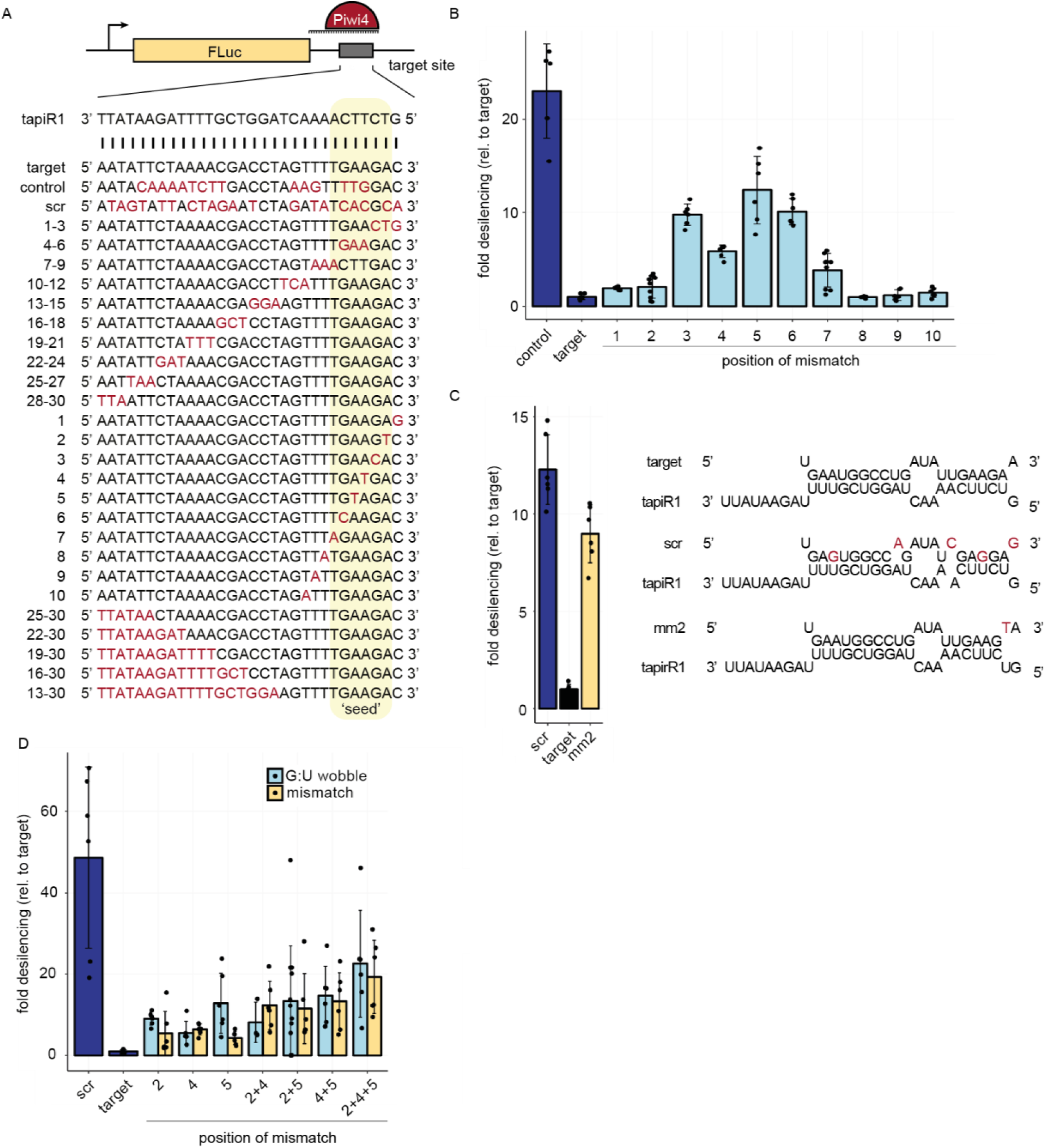
tapiR1 uses a G:U wobble sensitive seed sequence for target recognition. (A) Schematic representation of the reporter constructs used in panel B and Figure 2. Numbers indicate the position of the mismatch relative to the 5’ end of the piRNA. (B) Luciferase assay of reporters carrying a tapiR1 target site with single mismatches in the 3’ UTR. (C) Luciferase activity of reporters with the tapiR1 target site from RLuc and indicated mismatches in the 3’ UTR of FLuc (left panel). The tapiR1 target duplexes and mutants are presented in the right panel. (D) Luciferase activity of tapiR1 reporters carrying mismatches or G:U wobble base pairs at the indicated positions. Firefly luciferase activity was normalized to the activity of a co-transfected *Renilla* luciferase reporter to control for differences in transfection efficiencies. Data represent mean, standard deviation and individual measurements of representative experiments with two independent clones per construct and measured in triplicates.

**Extended Data Figure 7:**
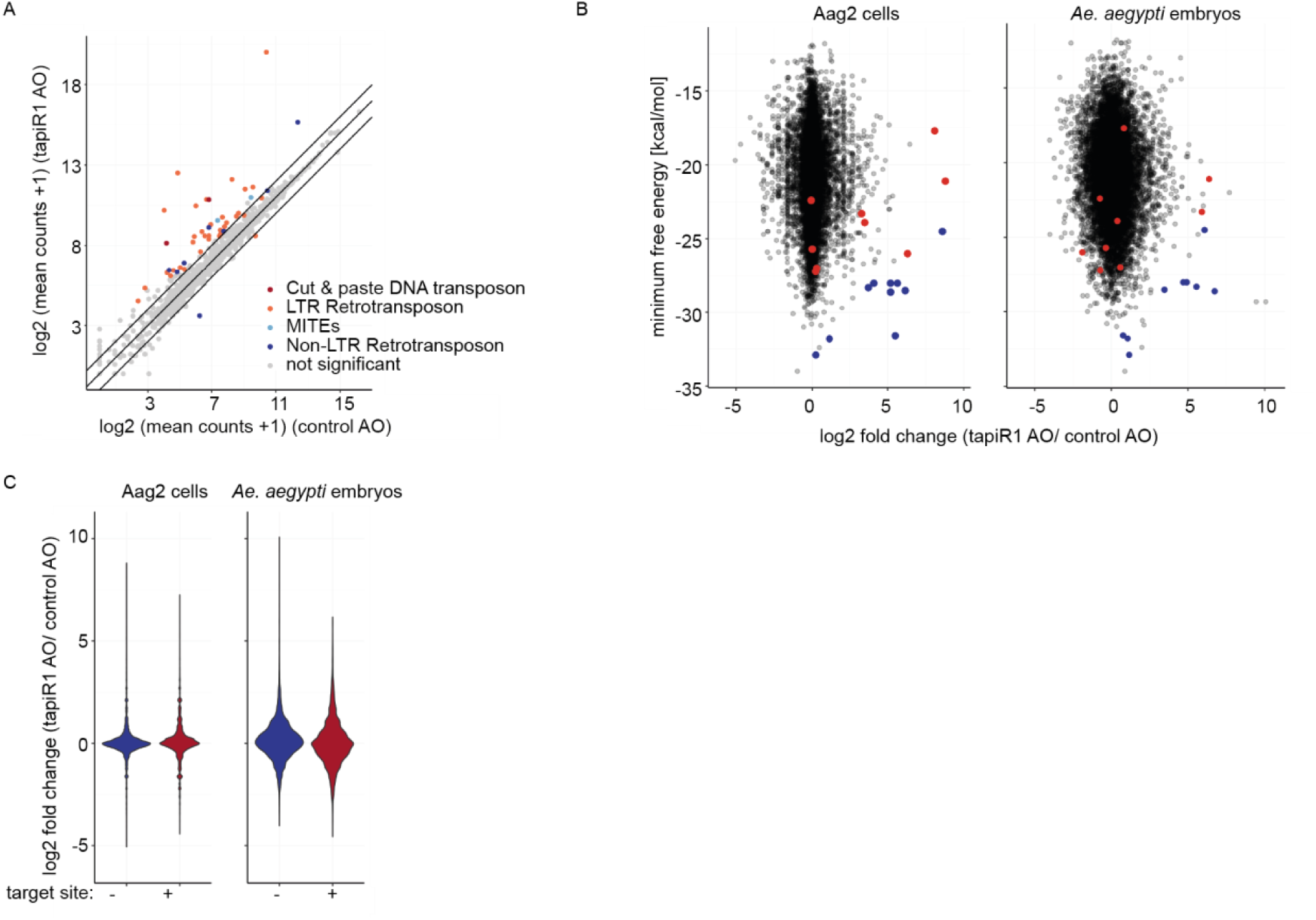
tapiR silences gene expression in Aag2 cells. (A) log2 mRNA expression of transposable elements in Aag2 cells treated with a tapiR1 specific antisense oligonucleotide (AO) or control AO. Depicted are the means of three biological replicates. A pseudo-count of one was added to all values in order to plot values of zero. Diagonal lines represent a fold change of two. Significance was tested at an FDR of 0.01 and a log2 fold change of 0.5. (B) log2 fold changes of genes upon treatment with tapiR1 or control AO in Aag2 cells (left) and mosquito embryos (right) plotted against the minimum free energy of predicted tapiR1-target duplexes. Blue dots indicate target sites that were confirmed to be functional, and red dots indicate target sites that were not functional in luciferase reporter assays (see Extended Data Fig 8). (C) Violin plot of log2 fold changes of all genes in Aag2 cells (left) and mosquito embryos (right), either with or without predicted tapiR1 target site.

**Extended Data Figure 8:**
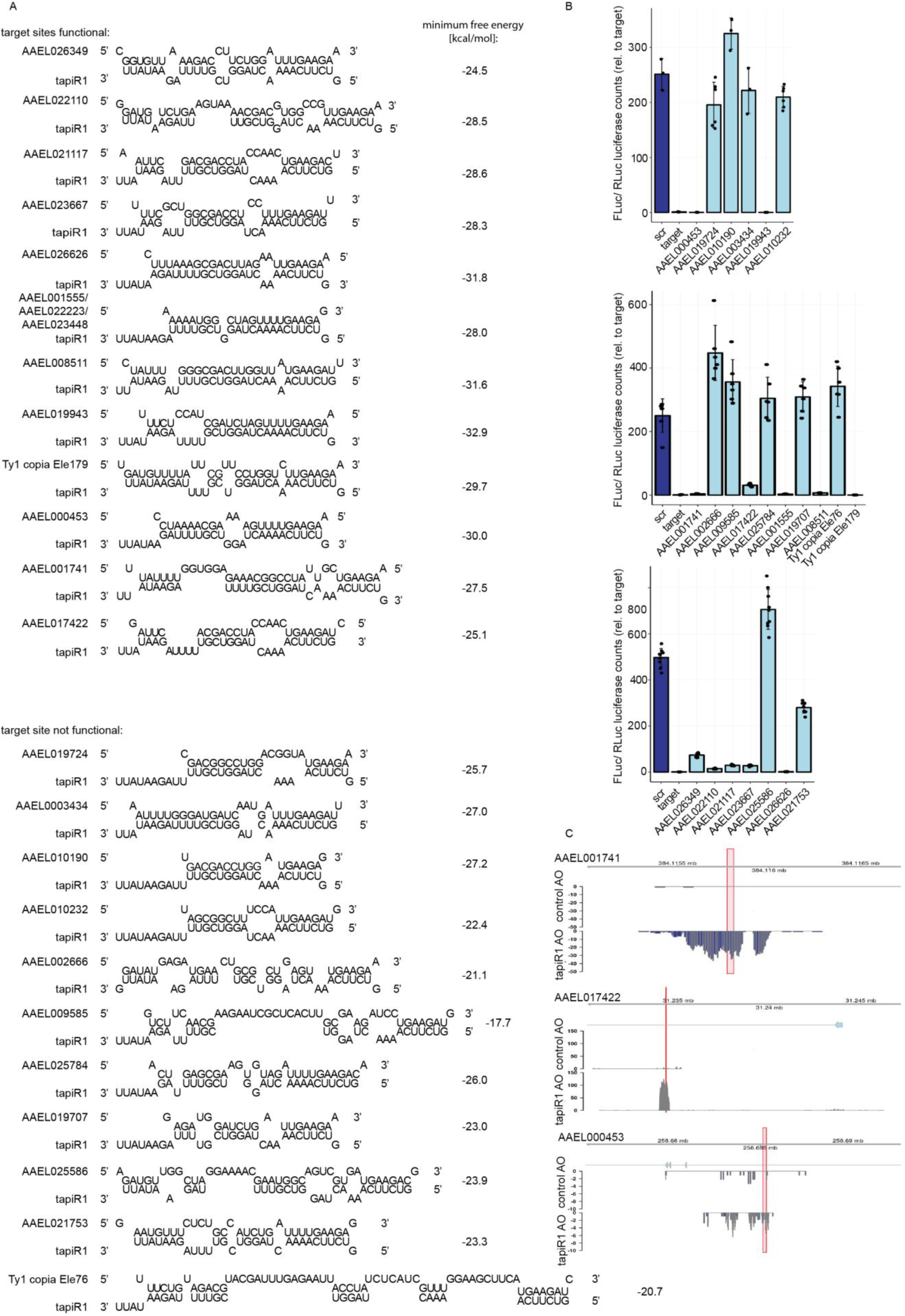
Validation of tapiR1 target genes. (A) Predicted structures and minimum free energy of tapiR1/target duplexes analysed in panel B. (B) Luciferase assay of reporters carrying the predicted target site from panel A in the 3’ UTR of firefly luciferase. Firefly luciferase activity was normalized to the activity of a co-transfected *Renilla* luciferase reporter to control for differences in transfection efficiencies. Indicated are mean, standard deviation and individual measurements from representative experiments performed with two to three independent clones per construct and measured in triplicates. (C) AAEL017422, AAEL001741, and AAEL000453 were annotated in the previous AaegL3 gene set, but not in the current AaegL5 gene set. Read coverage in tapiR1 AO and control AO treated Aag2 cells at these genomic regions suggests that these regions are actively transcribed, but repressed by tapiR1. Red boxes indicate the positions of tapiR1 target sites.

**Extended Data Figure 9:**
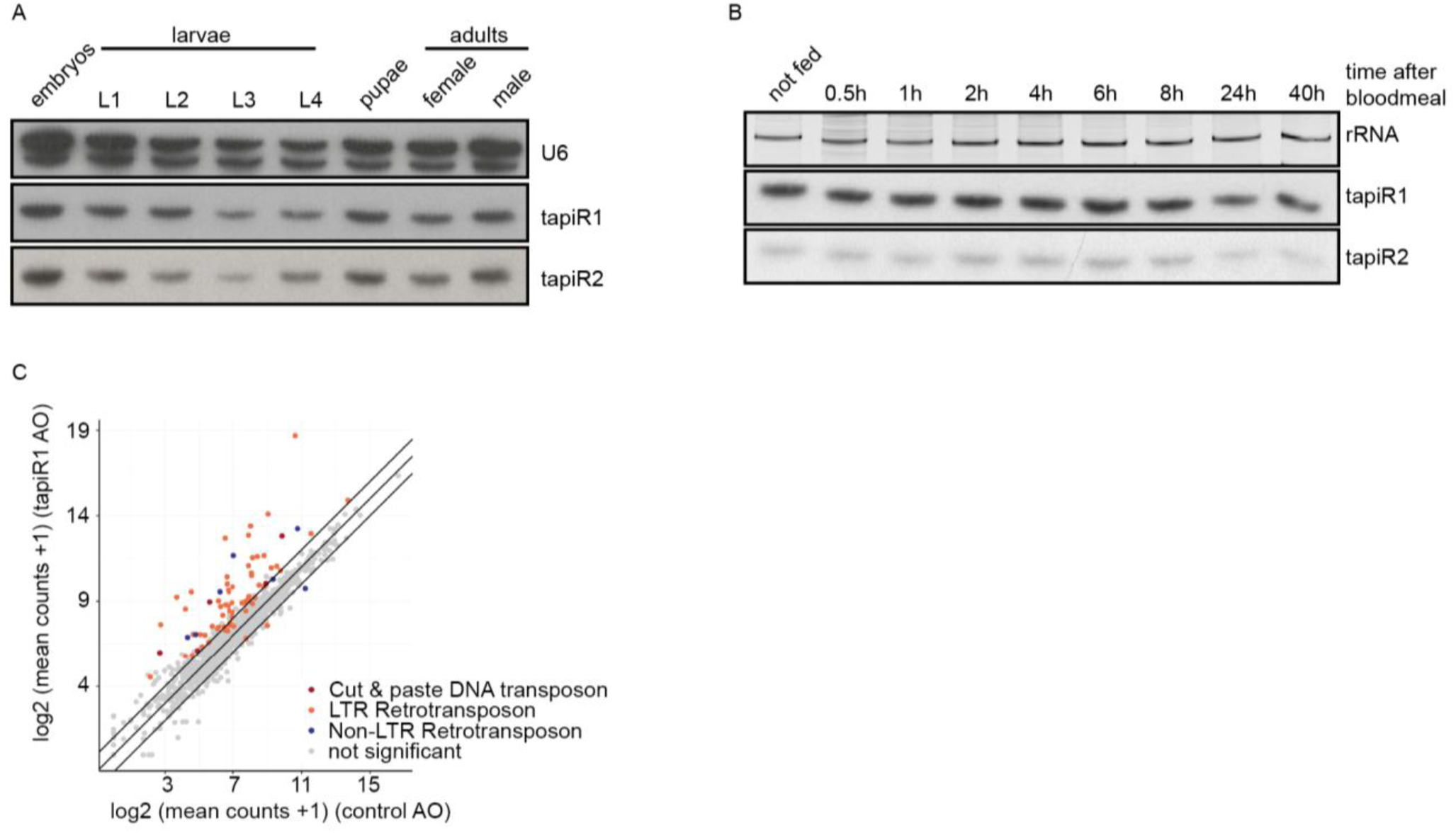
tapiR1 regulates gene expression in mosquito embryos. (A, B) Northern blot analysis of tapiR1 and 2 in developmental stages of *Ae. aegypti* mosquitoes (A), or at different time points after blood feeding (B). U6 snRNA (A) or ethidium bromide-stained rRNA (B) were analyzed to verify equal loading. (C) log2 mRNA expression of transposable elements in embryos injected with tapiR1-specific or control AO. Mean counts of five biological replicates are shown. Significance was tested at an FDR of 0.01 and a log2 fold change of 0.5. Diagonal lines indicate a fold change of two.

**Supplementary Data Figure S1:**
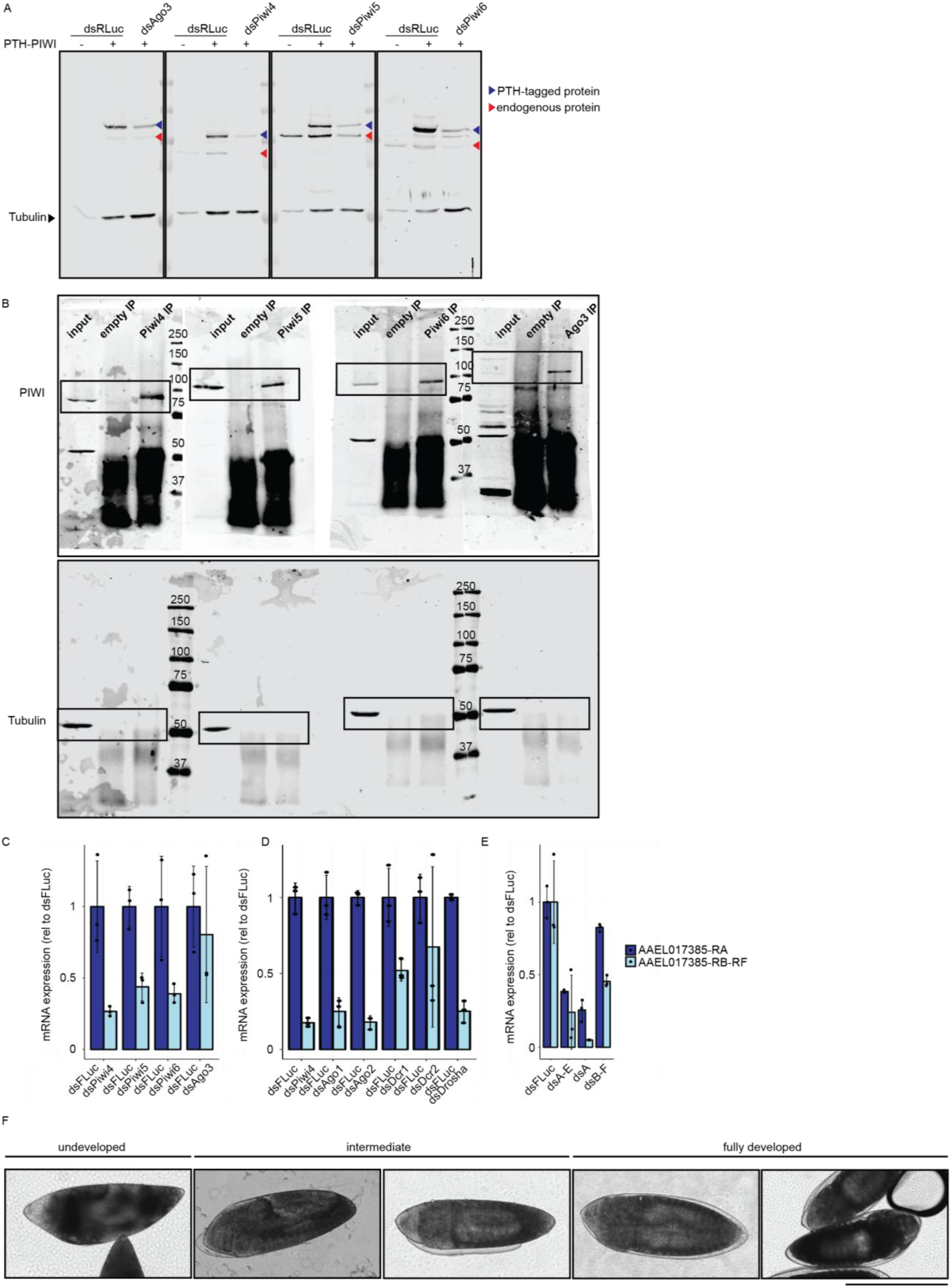
Antibody validation, uncropped Western blot images, knockdown efficiencies, and scoring scheme for the development of *Ae. aegypti* embryos. (A) Validation of *Ae. aegypti* PIWI antibodies. Specificity was confirmed by detection of an additional band in PTH-tagged PIWI-expressing Aag2 cells, and loss of signal upon dsRNA-mediated knockdown. Knockdown with dsRNA targeting RLuc (dsRLuc) serves as negative control. (B) Uncropped Western blot images corresponding to Extended Data Fig 2B. (C-E) Knockdown efficiencies of PIWI genes shown in Extended Data Fig 2D (D), siRNA and miRNA pathway genes shown in Extended Data Fig 2E (E), and AAEL017385 isoforms in the experiment shown in Extended Data Fig 3C (F) (F) Representative images of embryos scored as either undeveloped, intermediate or fully developed at 2.5 days post injection with antisense RNA oligonucleotides.

